# Chromosome-level genome assembly of *Helichrysum odoratissimum*, a medicinal plant from Southern Africa

**DOI:** 10.64898/2026.02.23.707396

**Authors:** Ansia van Coller, Victoria Ingle Cole, Sadik Muzemil, Samira Ghoor, Enrico Carlo Roode, Nadia Carstens, Brigitte Glanzmann, Renée Prins, Julian Onyewuonyeoma Osuji, Gane Ka-Shu Wong, Xun Xu, ThankGod Echezona Ebenezer, Craig John Kinnear

## Abstract

*Helichrysum odoratissimum*, a plant species native to Southern Africa, holds deep cultural significance and is widely used in traditional medicine. It is valued for its antimicrobial and anti-inflammatory properties and is cultivated and traded in South Africa due to its growing commercial relevance to the pharmaceutical and cosmetic industries. Despite its ecological and economic importance, genomic resources for this species remain limited.

Here, we report a high-quality chromosome-level genome assembly for *H. odoratissimum* to facilitate investigations into its genetic basis of bioactivity, environmental adaptation, and taxonomic relationships. High-molecular-weight DNA extracted from leaf tissue was sequenced using Oxford Nanopore Technologies (ONT) long-read sequencing, and Hi-C proximity ligation data were generated to enable chromosome-scale scaffolding. The final assembly spans 2.4 Gb, with a scaffold N50 of 354 Mb, and was anchored and oriented into seven chromosomes, with 2,935 smaller scaffolds remaining unplaced. Assembly quality and completeness were evaluated using QUAST and BUSCO, and chromosomal structure was validated using Hi-C contact maps.

This genome provides a foundational resource for functional genomics, conservation planning, molecular breeding, and the discovery of bioactive compounds underpinning the medicinal and commercial value of *H. odoratissimum*.

## Background & Summary

The genus *Helichrysum* of the Asteraceae family comprises over 600 species distributed across Africa, Asia, Australia, Madagascar and Europe, with a substantial portion being native to Southern Africa^1^. They are found in a range of habitats, often thriving in sub-montane regions, and their broad geographic range has promoted diversification into ecotypes and distinct taxa occupying different ecotunes and ecozones^2,3^. *Helichrysum odoratissimum* (Figure 1A), commonly known as “the everlasting plant” or *imphepho*, is native to Southern Africa and is adapted to warmer, more humid regions because of its climatic niche conservatism^4^. Although it is not currently considered threatened^5^, limited investment in its genetic improvement and genomics highlights the need for sustainable harvesting strategies and a clearer understanding of its adaptive potential in different climates. Importantly, ecological diversification within *Helichrysum* has been accompanied by extensive chemical variation, suggesting that genomic architecture may underlie both environmental resilience and bioactive compound production. However, the genetic basis of these traits in *H. odoratissimum* remains largely unexplored.

**Figure 1.**
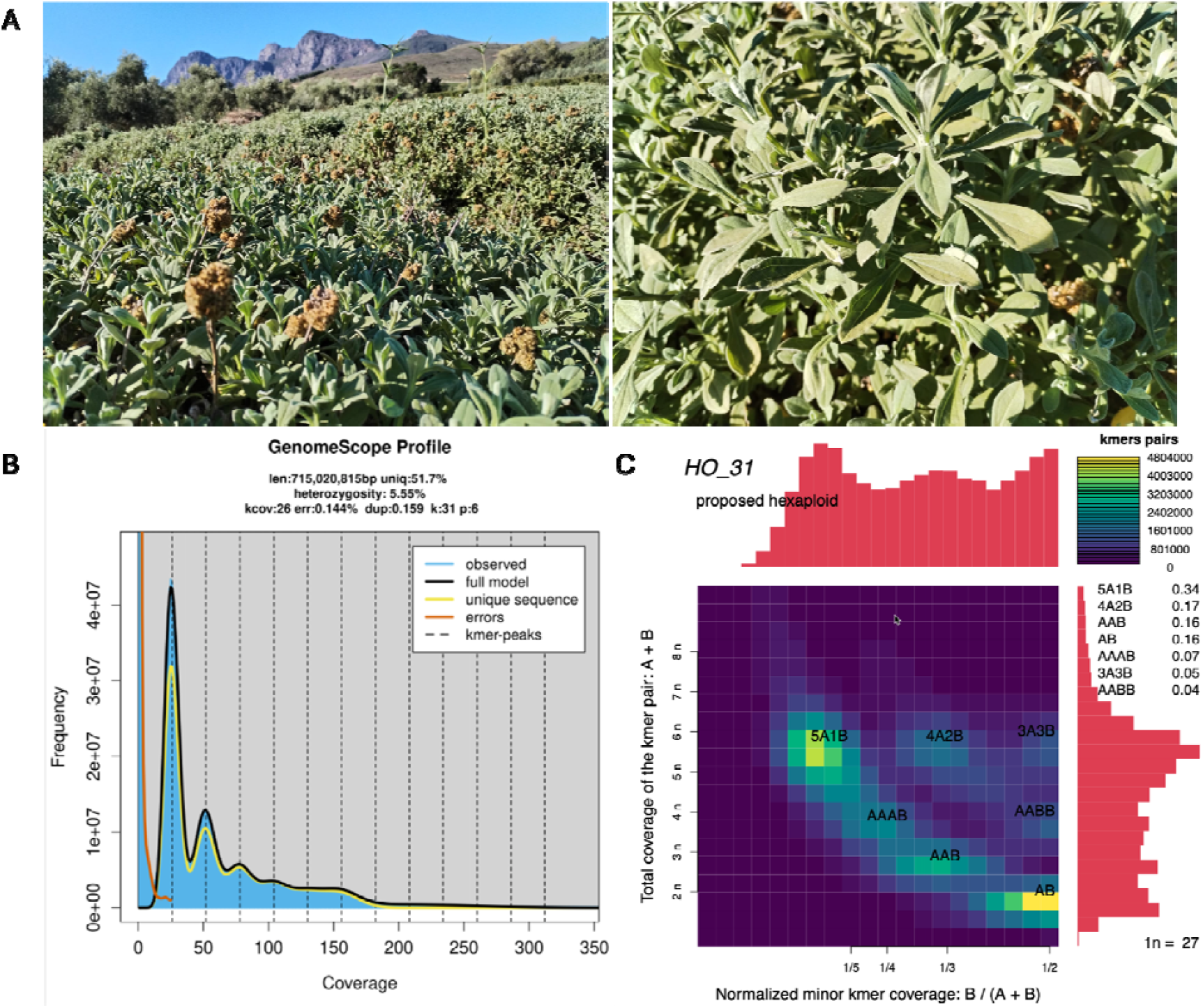
*Helichrysum odoratissimum* and estimates of genome size and ploidy level. (A) *H. odoratissimum* at Babylonstoren Estate in Cape Town, South Africa, where it was harvested. (B) GenomeScope plot of kmer size 31 (k=31) and ploidy level 6n, and (C) Smudgeplot (k=31) strong signal for 6n, suggesting haploid genome size of 715 MB and hexaploidy state of *H. odoratissimum*.

In South Africa, *H. odoratissimum* is widely harvested and traded for traditional medicine and ritual purposes, reflecting its rich phytochemical profile. Is cultivated for herbal teas, essential oils and aromatic insect-repellents^6–9^. Ethnomedicinal applications span gastrointestinal, respiratory, inflammatory, dermatological, and metabolic conditions, including liver disorders, stomach pain, colds, coughs, skin infections, hypertension, arthritis, and diabetes mellitus^7,10–14^. Moreover, extracts of *H. odoratissimum* exhibit antibacterial activity against *Staphylococcus, Enterococcus* and *Streptococcus* species^15,16^, antifungal activity against *Botrytis cinerea*^17^, inhibit quorum sensing in *Chromobacterium violaceum*^18^ and display anti-HIV-1 activity and CD4-boosting properties in cell-based assays^19–21^. Beyond therapeutic applications, the species shows commercial promise due to its nutritional and antioxidant properties. Its leaves and stems are rich in phytate, vitamins B_2_, C, and E, and minerals such as potassium, calcium, sodium, magnesium, phosphorus, and iron^22^. Extracts are being investigated as natural ingredients for pharmaceutical and cosmetic formulations, where they provide antioxidant photoprotection, enhance sunscreen photostability without mutagenicity, and demonstrate cytotoxic activity against human epidermoid carcinoma cells^23,24^. Understanding the genetic basis of this biochemical diversity is essential for both conservation and targeted bioprospecting.

Cytogenetic and molecular studies in *Helichrysum* species have revealed substantial genomic diversity. Chromosome number and ploidy levels vary widely across the genus^25^, and genome sizes range from 8.13 pg (2C) in H. rubicundum to 18.4 pg in H. leucocephalum and H. davisianum^26^. Population genetic studies employing Amplified Fragment Length Polymorphism (AFLP) and Inter Simple Sequence Repeat (ISSR) markers have further demonstrated pronounced genetic structuring and adaptive differentiation^2,3,27,28^. Multiple levels of ploidy have been reported within a single *Helichrysum* species suggesting that polyploidy is an important evolutionary driver for the diversification in the genus^29^. This chromosomal diversity likely contributes to the broad ecological adaptability and chemical diversity observed in *Helichrysum* species.

Despite wide utilisation *Helichrysum* for local medicinal purpose, and its emerging potential for commercialisation in pharmaceutical and cosmetic industries, genomic resources remain scarce: among the 600 known species of *Helichrysum*, only *H. umbraculigerum* has a chromosome-level reference genome assembly^30^, limiting comparitve and functional genomics.

Here, we present the first chromosome-level genome assembly for *H. odoratissimum*, generated using Oxford Nanopore Technologies (ONT) long-read sequencing in combination with High-throughput Chromosome Conformation Capture (Hi-C) scaffolding. To our knowledge, this represents the first chromosome-scale plant genome assembly done entirely on African soil using integrated ONT and MGI-Tech Hi-C platforms. As of 2022, of the 798 plant species sequenced globally, only ∼20 are endemic to Africa, and none had been assembled to chromosome scale on the continent^31,32^. In two previous studies led by African institutions, ONT sequencing was performed locally, but samples were exported for Hi-C processing^33,34^. The near-complete genome reported here provides a foundational resource for functional genomics, molecular breeding, and conservation planning, and establishes a platform for dissecting the genetic basis of medicinal traits and adaptive evolution within *Helichrysum*.

## Methods

### Ethics statement

The sample collection procedure, including the processing and handling of the plant material used in this study, complied with applicable institutional, national, and international regulations governing access to biological resources. Plant material of *H. odoratissimum* was obtained from an *ex situ* specimen, with written permission from the landowner. A collection and sampling permit was issued under the provisions of the Nature Conservation Ordinance, 1974, by CapeNature (permit reference number: CN35-28-35536), the provincial conservation authority responsible for biodiversity management in the Western Cape of South Africa.

Given the documented medicinal use of *Helichrysum* species, the planned research activities were classified under bioprospecting regulations, and the South African Department of Forestry, Fisheries and the Environment (DFFE) was formally notified of the study in accordance with the National Environmental Management: Biodiversity Act (NEMBA), 2004, and its associated Bioprospecting, Access and Benefit-Sharing (BABS) Regulations.

All procedures were conducted in alignment with ethical best practices for plant genomics research and guided by the African BioGenome Project (AfricaBP) Ethics, Legal, and Social Issues practical guide on accessing and sharing biological materials^35^ including compliance with access and benefit-sharing principles under the Convention on Biological Diversity and the Nagoya Protocol. As this work was undertaken solely for non-commercial academic research purposes, consultations with the relevant authorities confirmed that no immediate benefit-sharing agreement was required beyond permit compliance and reporting obligations. Any future downstream use with potential commercial application would be subject to additional regulatory approval and the establishment of formal benefit-sharing arrangements.

#### Plant materials, nucleic acid extraction and sequencing

*H. odorotissimum* stems and leaves were sampled at Babylonstoren estate in Cape Town, South Africa. Fresh leaves harvested from this mature plant grown under natural conditions were used to extract genomic DNA for ONT long-read sequencing. DNA was extracted using the Qiagen DNeasy plant pro kit (Qiagen, Hilden, Germany). RNA was extracted using the Qiagen RNeasy PowerPlant Kit (Qiagen, Hilden, Germany). DNA and RNA purity, yield and integrity were assessed using the Multiskan SkyHigh Microplate Spectrophotometer (ThermoFisher Scientific, MA, USA), Qubit 4 Fluorometer (ThermoFisher Scientific, MA, USA) and TapeStation 4200 System (Agilent Technologies, CA, USA).

For ONT long-read sequencing, 2µg high molecular weight (>60kb) genomic DNA was sheared to ∼13kb fragments using the Covaris G-Tube (PerkinElmer, MA, USA) and centrifugation at 4200 rpm for 60 seconds. Libraries were prepared using the ligation sequencing DNA kit (SQK-LSK114) (ONT, Oxford, UK). The library was loaded onto a R10.4.1 flow cell (FLO-PRO114M) and sequenced on the PromethION P2 Solo (ONT, Oxford, UK) for 102 hours. Sequencing was controlled using MinKNOW (v24.02.16), and base calling was performed using Dorado (v7.3.11) with the super-accurate (SUP) model.

Hi-C analysis was performed to capture genome-wide chromatin interactions in *H. odoratissimum* as previously described^36^. Briefly, healthy leaf tissue (∼0.4 g) was harvested and immediately cross-linked to preserve chromatin interactions. Tissue was finely ground under liquid nitrogen and cross-linked in 1% formaldehyde for 20 minutes at room temperature, quenched with 125 mM glycine, washed, pelleted, and stored at - 20 °C until nuclei isolation. Nuclei were extracted in chilled nuclear isolation buffer, permeabilized with SDS and Triton X-100, digested overnight with DpnII, end-filled with a biotinylated nucleotide mix, and proximity-ligated under dilute conditions; cross-links were then reversed, and DNA was purified and quantified. Finally, libraries were prepared using the MGIEasy Universal DNA Library Prep Kit (MGI Tech, Shenzhen,

China) and sequenced on the DNBSeq-G400 (MGI Tech, Shenzhen, China) for 64 hours. A paired-end sequencing strategy was employed, with a read length of 150bp. RNA libraries were prepared using the MGIEasy RNA Library Prep Kit (MGI Tech, Shenzhen, China) and sequenced on the DNBSeq-G400. In total, we obtained 140 Gb of ONT data, 73 Gb of Hi-C data, and 80 Gb of RNA-seq data.

#### Data pre-processing

The quality of ONT reads was assessed using NanoPlot (v1.44.0)^37^. Sequencing adapters were trimmed using Porechop (v0.2.4)^38^, and Chopper (v0.8.0)^37^ was used to filter out low base quality score (<Q10) and short reads (<1kb). Hi-C read quality was assessed using FastQC (v0.12.0)^39^ and MultiQC (v1.25.2)^40^, and Trimmomatic (v0.40.)^41^ was used to remove low-quality bases (<Q20) and trim the first 10bp from each read.

#### Estimation of genome size

Error-corrected ONT reads were used to estimate genome size and ploidy through k-mer analysis. KMC (v3.2.4)^42^ was used to generate k-mer histograms for k-mer size of 31. GenomeScope (v2.0)^43^ estimated the haploid genome size to be approximately 715 Mb. Ploidy estimation using Smudgeplot (v0.4.0)^43^ indicated that this *H. odoratissimum* individual is likely hexaploid.

#### De novo assembly

De novo assembly was performed with hifiasm (v0.2.4)^44,45^ using parameters --ont, --telo-m TTTAGGG, and --n-hap 6. GFA files were converted to FASTA format using gfatools (v0.5) ^46^. Assembly completeness was assessed with QUAST (v5.2) ^47^ and Benchmarking Universal Single-Copy Orthologs (BUSCO) (v5.8.0) ^48^ against the eudicots_odb10 lineage.

To reduce organelle-derived contamination, contigs <50kb were aligned using BLASTn (v2.16.0+)^49^ against a custom database of mitochondrial and plastid genomes obtained from the NCBI RefSeq organelle genome collections (mitochondrial and plastid; accessed March 2025)^50^. Contigs with >50% sequence identity to organelle sequences were removed.

Hi-C reads were mapped to the filtered assemblies using BWA-MEM2 (v2.2.1) ^51^ with the -5SP flag. Pairtools (v1.1.0)^52^ was used to filter for valid Hi-C contacts. Scaffolding was then performed using YaHS (v1.2.2)^53^ with --telo-motif TTTAGGG. Hi-C data were re-mapped to the scaffolded assemblies to evaluate contact maps. PretextMap (v0.1.9) (https://github.com/sanger-tol/PretextMap) and PretextView (v0.2.5) (https://github.com/sanger-tol/PretextView) were used for manual curation, supported by the Sanger rapid-curation pipeline (https://github.com/sanger-tol/rapid-curation).

#### Genome curation

Hi-C contacts were viewed in PretextView in edit mode and various manual re-arrangements were made. In total 68 scaffolds, accounting for 96,4% of the genome, were anchored to 7 individual chromosomes (Figure 2A).

**Figure 2.**
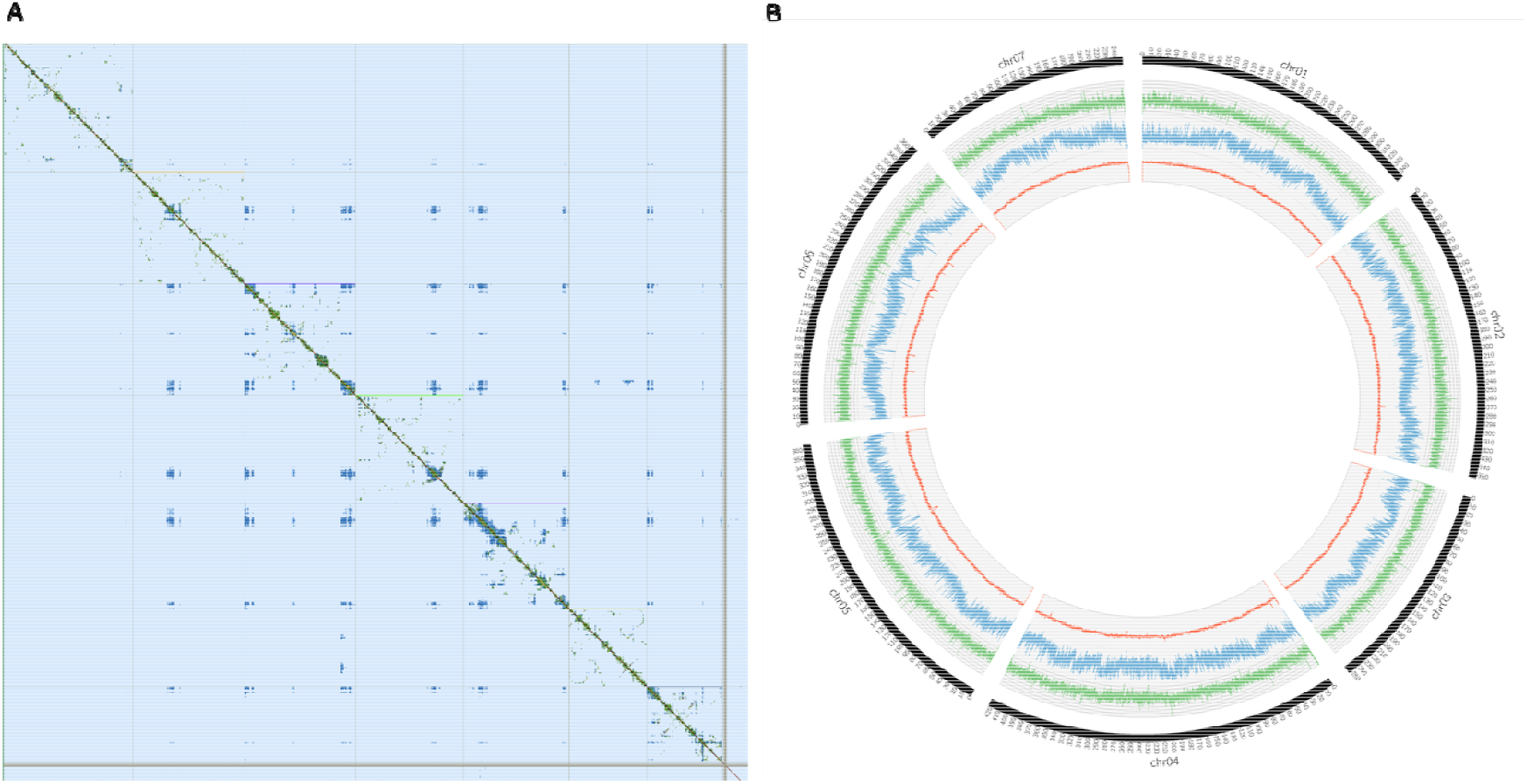
The assembled *Helichrysum odoratissimum* genome. (A) Hi-C interaction heatmap of the *H. odoratissimum* genome anchored to 7 large chromosomes. (B) Circos plot of the *H. odoratissimum* genome. From the outer ring to the inner ring, the tracks represent chromosome length, gene density, repeat density, and GC content.

#### Repeat and Gene Content

To identify repetitive elements, GMATA^54^ was used to identify Simple Sequence Repeats (SSRs), Tandem Repeat Finder (TRF)^55^ was used to identify Tandem repeats (TRs), and RepeatModeler and RepeatMasker^56^ were used to identify Transposable elements (TEs). BRAKER3^57^ was used for gene structure annotation in conjunction with RNA-seq data. To reduce redundancy and remove fragmented predictions, the initial BRAKER protein set was clustered at 95% identity using CD-HIT^58^ and proteins shorter than 150 amino acids were filtered out, retaining only high-confidence models. A total of 101,167 protein-coding genes were predicted (Figure 2B). The assembled genome contained 0.58% SSRs, 8,53% TRs and 53,9% TEs.

### Data Records

The raw sequencing data and the assembled genome of *H. odoratissimum* have been deposited in the European Nucleotide Archive (ENA) under the BioProject number PRJEB108529. The corresponding ONT, Hi-C raw, and RNA-seq data can be accessed via ENA accession numbers ERR16709387, ERR16709411, and ERR16709412. The assembled genome and annotation has ENA accession numbers GCA_980990915.1 and ERZ29109081.

### Technical Validation

To assess the quality, completeness, and structure of the *H. odoratissimum* genome assembly, we applied several validation tools and metrics.

#### Genome size and ploidy estimates

GenomeScope (v2.0)^43^, was used to profile the genome based on a k-mer size 31 frequency distribution for error-corrected ONT reads. The estimated haploid genome size is 715 Mb, with a heterozygosity of 5.55%. Smudgeplot^43^ (v0.4.0) analysis suggested a hexaploid genome structure, which informed the assembly parameters used with hifiasm (v0.2.4)^44,45^. Figures 1B and 1C show the GenomeScope and Smudgeplot visualisations.

#### Assembly quality and completeness

The primary assembly was evaluated using QUAST (v5.2)^47^ and BUSCO (v5.8.0)^48^. BUSCO analysis was performed using the eudicots_odb10 database, which contains 2326 single-copy orthologs. The primary draft assembly recovered 98% complete BUSCOs, including 19% as single-copy and 79% as duplicated, consistent with expectations for a collapsed hexaploid genome. After organelle filtering, 1961 contigs were removed due to high similarity to mitochondrial or plastid sequences, resulting in a filtered assembly length of 2.45 Gb.

#### Read alignment back to assembly

ONT reads were aligned back to the final scaffolded genome using minimap2 (v2.24), and the mapping rate was 99% and the coverage was 57X, suggesting a high degree of assembly accuracy and completeness. Hi-C reads were aligned with BWA-MEM2 (v2.2.1) ^51^, and valid Hi-C contacts were processed by Pairtools (v1.1.0)^52^.

#### Hi-C data quality and scaffolding validation

After mapping and deduplication using Pairtools, ∼473.8 million Hi-C read pairs were processed, of which 156.2 million were mapped and 77.8 million retained as non-duplicates. Approximately 73.7% of non-duplicate contacts were intra-chromosomal (cis), indicative of high-quality proximity ligation and effective chromatin capture. Furthermore, 3.2% of cis contacts spanned more than 10 kb, supporting successful scaffolding potential. All cis contacts fell within the expected convergence distance across strands, reinforcing the structural validity of the data. These contact patterns were further visualized using PretextView, confirming chromosome-scale. The contact maps revealed strong, diagonal chromosome-scale contact patterns with some minimal noise or mis-joins, indicative of successful contig ordering and orientation (Figure 2). The final assembly was anchored to 7 chromosomes, which is the estimated chromosome number for this species^29^. The final assembly size is 2.45Gb, with an N50 of 354 Mb, and recovered 97.1% of BUSCO genes (S:33.3%, D:63.8%) (Table 1). These assessments indicate that this is a high quality, near-complete genome.

**Table 1:** Assembly statistics for *Helichrysum odoratissimum* genome.

|  | *Helichrysum odoratissimum* |
| --- | --- |
| K-mer estimated haploid size; predicted ploidy level | 715 Mb; 6n |
| Assembled bases | 2.45 Gb |
| Number of chromosomes | 7 |
| Unplaced sequences | 3,5% |
| Contig N50 | 7 Mb |
| Scaffold N50 | 353 Mb |
| Repeats content | 63% |
| Predicted protein-coding gene count | 101167 |
| GC content | 35.7% |
| BUSCO completeness | C:97.1% [S:33.3%;D:63,8%]* |

\* Predictions are k-mer based and derived from GenomeScope and Smudgeplot analysis
\*\*C=Complete Benchmarking Universal Single-Copy Orthologs (BUSCO); S=Complete and single-copy BUSCO; D=Complete and duplicated BUSCO

## Code availability

All bioinformatics tools utilised for this manuscript have been mentioned in the text and are freely available.

## Acknowledgements

We would like to acknowledge Dr Ernst van Jaarsveld of Babylonstoren for his assistance with sample identification and collection. The authors acknowledge the training support provided for ONT sequencing by the 1000 Genomes South Africa (1KSA) project (www.1KSA.org.za) with support provided by the Department of Science, Technology and Innovation’s South African Research Infrastructure Roadmap through DIPLOMICS (www.diplomics.org.za). This work has benefited from the support and resources provided by the South African Medical Research Council (SAMRC), MGI, and the African BioGenome Project (AfricaBP).

## Author contributions

The work presented in this manuscript was carried out in collaboration with all authors. All authors made contributions to the conception or design of the work and interpretation of the data. RP coordinated sample collection logistics and transport to the SAMRC labs. Nucleic acid extractions and Hi-C library preparation was done by SG. ONT sequencing experiments were done by VIC. RNA and Hi-C sequencing was done by ER. Bioinformatics analysis was done primarily by AvC, with substantial input from SM, GKW, NC, BG, CJK. Manual curation of the genome was performed by SM. AvC, VIC, CJK, and TEE drafted the manuscript. AvC coordinated ethical compliance activities with input from CJK and TEE. CJK and TEE performed leadership responsibilities and project oversight. All authors critically revised and approved the final manuscript. All authors agreed to be accountable for all aspects of the work.

## References

1. Plants of Southern Africa: An Annotated Checklist. (National Botanical Institute, Pretoria, 2003).

2. Ninčević, T. et al. Population structure and adaptive variation of Helichrysum italicum (Roth) G. Don along eastern Adriatic temperature and precipitation gradient. Sci. Rep. 11, 24333 (2021).

3. Galbany-Casals, M., Blanco-Moreno, J. M., Garcia-Jacas, N., Breitwieser, I. & Smissen, R. D. Genetic variation in Mediterranean Helichrysum italicum (Asteraceae; Gnaphalieae): do disjunct populations of subsp. microphyllum have a common origin? Plant Biol. 13, 678–687 (2011).

4. Glennon, K. L. & Cron, G. V. Climate and leaf shape relationships in four Helichrysum species from the Eastern Mountain Region of South Africa. Evol. Ecol. 29, 657–678 (2015).

5. von Staden, L. Helichrysum odoratissimum (L.) Sweet. National Assessment: Red List of South African Plants version 2024.1. Accessed on 2026/02/17. (2010).

6. Mastelić, J., Politeo, O. & Jerković, I. Contribution to the Analysis of the Essential Oil of Helichrysum italicum (Roth) G. Don. – Determination of Ester Bonded Acids and Phenols. Molecules 13, 795–803 (2008).

7. Lourens, A. C. U., Viljoen, A. M. & van Heerden, F. R. South African Helichrysum species: A review of the traditional uses, biological activity and phytochemistry. J. Ethnopharmacol. 119, 630–652 (2008).

8. Abbas, M. G. et al. Chemical Composition, Repellent, and Oviposition Deterrent Potential of Wild Plant Essential Oils against Three Mosquito Species. Molecules 29, 2657 (2024).

9. Moteetee, A., Moffett, R. O. & Seleteng-Kose, L. A review of the ethnobotany of the Basotho of Lesotho and the Free State Province of South Africa (South Sotho). South Afr. J. Bot. 122, 21–56 (2019).

10. De Canha, M. N., Komarnytsky, S., Langhansova, L. & Lall, N. Exploring the Anti-Acne Potential of Impepho [Helichrysum odoratissimum (L.) Sweet] to Combat Cutibacterium acnes Virulence. Front. Pharmacol. 10, (2020).

11. de la Garza, A.L. et al. Helichrysum and Grapefruit Extracts Boost Weight Loss in Overweight Rats Reducing Inflammation. J. Med. Food 18, 890–898 (2015).

12. Erasto, P., Adebola, P. O., Grierson, D. S. & Afolayan, A. J. An ethnobotanical study of plants used for the treatment of diabetes in the Eastern Cape Province, South Africa. Afr. J. Biotechnol. 4, (2005).

13. Nortje, J. M. & van Wyk, B.-E. Medicinal plants of the Kamiesberg, Namaqualand, South Africa. J. Ethnopharmacol. 171, 205–222 (2015).

14. Olorunnisola, O. S., Bradley, G. & Afolayan, A. J. Ethnobotanical information on plants used for the management of cardiovascular diseases in Nkonkobe Municipality, South Africa. J. Med. Plants Res. 5, 4256–4260 (2011).

15. Mathekga, A. D. M. & Meyer, J. J. M. Antibacterial activity of South African Helichrysum species. South Afr. J. Bot. 64, 293–295 (1998).

16. Serabele, K. et al. Comparative chemical profiling and antimicrobial activity of two interchangeably used ‘Imphepho’ species (Helichrysum odoratissimum and Helichrysum petiolare). South Afr. J. Bot. 137, 117–132 (2021).

17. Matrose, N. A., Belay, Z. A., Obikeze, K., Mokwena, L. & Caleb, O. J. Bioprospecting of Helichrysum Species: Chemical Profile, Phytochemical Properties, and Antifungal Efficacy against Botrytis cinerea. Plants 12, 58 (2023).

18. Moyo, P. et al. Quorum sensing inhibition by South African medicinal plants species: an in vitro and an untargeted metabolomics study. BMC Complement. Med. Ther. 25, 138 (2025).

19. Heyman, H. M. et al. Identification of anti-HIV active dicaffeoylquinic- and tricaffeoylquinic acids in Helichrysum populifolium by NMR-based metabolomic guided fractionation. Fitoterapia 103, 155–164 (2015).

20. Salehi, B. et al. Medicinal Plants Used in the Treatment of Human Immunodeficiency Virus. Int. J. Mol. Sci. 19, 1459 (2018).

21. Yazdi, S. E. et al. Anti-HIV-1 activity of quinic acid isolated from Helichrysum mimetes using NMR-based metabolomics and computational analysis. South Afr. J. Bot. 126, 328–339 (2019).

22. Afuape, A. O., Afolayan, A. J. & Buwa-Komoreng, L. V. Proximate, Vitamins, Minerals and Anti-Nutritive Constituents of the Leaf and Stem of Helichrysum odoratissimum (L.) Sweet: A Folk Medicinal Plant in South Africa. Int. J. Plant Biol. 13, 463–472 (2022).

23. Twilley, D. et al. The Effect of Helichrysum odoratissimum (L.) Sweet on Cancer Cell Proliferation and Cytokine Production. Int. J. Pharmacogn. Phytochem. Res. 9, 621– 631 (2017).

24. Twilley, D. et al. Ethanolic extracts of South African plants, Buddleja saligna Willd. and Helichrysum odoratissimum (L.) Sweet, as multifunctional ingredients in sunscreen formulations. South Afr. J. Bot. 137, 171–182 (2021).

25. Glennon, K. L. & Cron, G. V. DNA Ploidy Variation and Population Structure of the Morphologically Variable Helichrysum odoratissimum (L.) Sweet (Asteraceae) in South Africa. Int. J. Plant Sci. 180, (2019).

26. Azizi, N., Sheidai, M., Mozaffarian, V. & Nourmohammadi, Z. Karyotype and genome size analyses in species of Helichrysum (Asteraceae). Acta Bot. Bras. 28, 367–375 (2014).

27. Galbany-Casals, M., Carnicero-Campmany, P., Blanco-Moreno, J. M. & Smissen, R. D. Morphological and genetic evidence of contemporary intersectional hybridisation in Mediterranean Helichrysum (Asteraceae, Gnaphalieae). Plant Biol. 14, 789–800 (2012).

28. Melito, S. et al. Genetic and Metabolite Diversity of Sardinian Populations of Helichrysum italicum. PLOS ONE 8, e79043 (2013).

29. Galbany-Casals, M. & Romo, À.M. Polyploidy and new chromosome counts in Helichrysum (Asteraceae, Gnaphalieae). Bot. J. Linn. Soc. 158, 511–521 (2008).

30. Berman, P. et al. Parallel evolution of cannabinoid biosynthesis. Nat. Plants 9, 817– 831 (2023).

31. Marks, R. A., Hotaling, S., Frandsen, P. B. & VanBuren, R. Representation and participation across 20 years of plant genome sequencing. Nat. Plants 7, 1571– 1578 (2021).

32. Ebenezer, T. E. et al. Africa: sequence 100,000 species to safeguard biodiversity. Nature 603, 388–392 (2022).

33. Njaci, I. et al. Chromosome-level genome assembly and population genomic resource to accelerate orphan crop lablab breeding. Nat. Commun. 14, 1915 (2023).

34. Waweru, B. et al. Chromosome-scale assembly of the African yam bean genome. Sci. Data 11, 1384 (2024).

35. AfricaBP ELSI Subcomittee. Practical Guide to Accessing and Sharing Biological Diversity Material and Data: A Research Stage-Based Approach. https://doi.org/10.17605/OSF.IO/83DXN (2025) doi:10.17605/OSF.IO/83DXN.

36. Lieberman-Aiden, E. et al. Comprehensive mapping of long-range interactions reveals folding principles of the human genome. Science 326, 289–293 (2009).

37. De Coster, W. & Rademakers, R. NanoPack2: population-scale evaluation of long-read sequencing data. Bioinformatics 39, btad311 (2023).

38. Bonenfant, Q., Noé, L. & Touzet, H. Porechop_ABI: discovering unknown adapters in Oxford Nanopore Technology sequencing reads for downstream trimming. Bioinforma. Adv. 3, vbac085 (2023).

39. Andrews, S. FastQC A Quality Control tool for High Throughput Sequence Data. (2010).

40. Ewels, P., Magnusson, M., Lundin, S. & Käller, M. MultiQC: summarize analysis results for multiple tools and samples in a single report. Bioinformatics 32, 3047– 3048 (2016).

41. Bolger, A. M., Lohse, M. & Usadel, B. Trimmomatic: a flexible trimmer for Illumina sequence data. Bioinformatics 30, 2114–2120 (2014).

42. Kokot, M., Długosz, M. & Deorowicz, S. KMC 3: counting and manipulating k-mer statistics. Bioinformatics 33, 2759–2761 (2017).

43. Ranallo-Benavidez, T. R., Jaron, K. S. & Schatz, M. C. GenomeScope 2.0 and Smudgeplot for reference-free profiling of polyploid genomes. Nat. Commun. 11, 1432 (2020).

44. Cheng, H., Asri, M., Lucas, J., Koren, S. & Li, H. Scalable telomere-to-telomere assembly for diploid and polyploid genomes with double graph. Nat. Methods 21, 967–970 (2024).

45. Cheng, H. et al. Efficient near telomere-to-telomere assembly of Nanopore Simplex reads. 2025.04.14.648685 Preprint at 10.1101/2025.04.14.648685 (2025).

46. Pani, S., Dabbaghie, F., Marschall, T. & Söylev, A. gaftools: a toolkit for analyzing and manipulating pangenome alignments. 2024.12.10.627813 Preprint at 10.1101/2024.12.10.627813 (2024).

47. Gurevich, A., Saveliev, V., Vyahhi, N. & Tesler, G. QUAST: quality assessment tool for genome assemblies. Bioinformatics 29, 1072–1075 (2013).

48. Seppey, M., Manni, M. & Zdobnov, E. M. BUSCO: Assessing Genome Assembly and Annotation Completeness. Methods Mol. Biol. 1962, 227–245 (2019).

49. Camacho, C. et al. BLAST+: architecture and applications. BMC Bioinformatics 10, 421 (2009).

50. Sayers, E. W. et al. Database resources of the National Center for Biotechnology Information in 2025. Nucleic Acids Res. 53, D20–D29 (2025).

51. Vasimuddin, Md., Misra, S., Li, H. & Aluru, S. Efficient Architecture-Aware Acceleration of BWA-MEM for Multicore Systems. in 2019 IEEE International Parallel and Distributed Processing Symposium (IPDPS) 314–324 (2019). doi:10.1109/IPDPS.2019.00041.

52. Open2C et al. Pairtools: from sequencing data to chromosome contacts. 2023.02.13.528389 Preprint at 10.1101/2023.02.13.528389 (2023).

53. Zhou, C., McCarthy, S. A. & Durbin, R. YaHS: yet another Hi-C scaffolding tool. Bioinformatics 39, btac808 (2023).

54. Wang, X. & Wang, L. GMATA: An Integrated Software Package for Genome-Scale SSR Mining, Marker Development and Viewing. Front. Plant Sci. 7, (2016).

55. Benson, G. Tandem repeats finder: a program to analyze DNA sequences. Nucleic Acids Res. 27, 573–580 (1999).

56. Flynn, J. M. et al. RepeatModeler2 for automated genomic discovery of transposable element families. Proc. Natl. Acad. Sci. 117, 9451–9457 (2020).

57. Gabriel, L. et al. BRAKER3: Fully automated genome annotation using RNA-seq and protein evidence with GeneMark-ETP, AUGUSTUS and TSEBRA. BioRxiv Prepr. Serv. Biol. 2023.06.10.544449 (2024) doi:10.1101/2023.06.10.544449.

58. Li, W. & Godzik, A. Cd-hit: a fast program for clustering and comparing large sets of protein or nucleotide sequences. Bioinformatics 22, 1658–1659 (2006).

